# A noradrenergic I_h_-dependent pacemaker system drives sniffing

**DOI:** 10.64898/2026.05.28.728179

**Authors:** Babita Thadari, Youjin Lee, Jordan Skach, Marcelo Camba Almazán, Eun Jung Hwang, Kaiwen Kam

## Abstract

Sniffs are rapid breaths generated to clear nasal passages, actively sample air for smelling, and, in infants, rouse from sleep. Rapid shifts between breathing at rest (eupnea) and sniffing point to generation of both behaviors by the core brainstem inspiratory oscillator, the preBötzinger Complex (preBötC). Eupneic rhythm generating mechanisms, when driven faster, are assumed to underlie higher frequency sniffing rhythms; however, the mechanisms governing sniffing have not been directly examined and remain unknown. Using head-fixed awake mice and rhythmically active medullary slices containing preBötC, we find that norepinephrine in preBötC elicits sniff bouts whose hyperpolarization-activated cyclic nucleotide-gated (HCN/I_h_) channel-dependent rhythmogenic mechanisms are distinct from eupneic rhythm. These findings reveal how neuromodulation enables a single microcircuit to flexibly switch between distinct oscillatory modes, providing a mechanistic framework for rapid respiratory-related behavioral transitions and potential therapeutic targets for conditions affecting arousal and breathing.

## Introduction

Breathing at rest is optimized for efficient gas exchange to sustain life, but humans and other mammals mix regular breaths with sniffs, sighs, and other inspiratory patterns to maintain airway and lung clearance, rouse themselves, and smell^1-6^. Sniffing, in particular, expands respiratory functionality beyond airflow. Commonly associated with smelling, sniffs are short bouts of two or more breaths that occur at frequencies two to five times faster than breathing at rest (also called eupnea). Humans sniff at ∼1 Hz, compared to a eupneic frequency of ∼0.2-0.3 Hz, and mice sniff from 4-13 Hz, compared to a eupneic frequency of ∼2-4 Hz, slightly faster than rats^7-10^. Olfactory sniffs drive other orofacial behaviors and generate neural oscillations throughout the brain to modulate learning, memory, and emotion^11-14^. Beyond olfaction, disruption of higher frequency, sniff-like inspirations in infancy may prevent airway clearance, increasing risk of infection, and underlie the inability to arouse to breathe in Congenital Central Hypoventilation Syndrome (CCHS) and Sudden Infant Death Syndrome (SIDS)^4,15^,16. Abnormally rapid breathing at sniff-like frequencies is observed in panic or anxiety disorders^17^. Thus, higher frequency, sniff-like breathing sits at the core of a suite of arousal, active sensing, and exploratory behaviors, whose sensorimotor circuits are fundamental for brain function and even survival.

While the sensory arm of the olfactory sensorimotor circuit is well-characterized, much less is known about mechanisms producing and shaping sniffing motor behavior. While several brain regions, such as ventral striatum^18^ and parabrachial nucleus^19^, regulate sniffing, inspiratory patterns, including sniffing, are ultimately driven by the preBötzinger Complex (preBötC), a nucleus in the ventrolateral medulla identified in a number of mammalian species, including humans^11,20-22^. preBötC is necessary for generating eupneic inspiratory rhythm and seamlessly intermixes eupneic breaths with sighs and sniffs to meet physiological, emotive, and volitional needs^5,11^,20,22,23. Rapid switching between eupnea and sniffing, the resemblance of sniffs to eupneic breaths, and the tunability of breathing frequency have led to the assumption that the same rhythmogenic mechanisms underlie eupneic breathing and sniffing and that sniffing is produced by simply increasing preBötC excitability to drive rhythm faster.

Eupneic inspiratory rhythmogenesis is hypothesized to be an emergent process involving synchronization of developing brain homeobox protein 1 – derived (Dbx1^+^) glutamatergic interneurons^11,24,25^. Synchronization across the preBötC Dbx1^+^ network is thought to initiate the bursting phase by activating persistent inward conductances, such as Ca^2+^ -activated non-specific cation current (I_CAN_) and persistent sodium current (I_NaP_)^26-28^. Yet, increasing preBötC excitatory drive by altering extracellular K^+^ or Ca^2+^, adding excitatory neuromodulators *in vitro*, or even direct optogenetic stimulation of preBötC subpopulations, such as Dbx1^+^ neurons, *in vivo*, fail to produce sniff-like frequencies^29-31^. Thus, despite the importance of sniffs for arousal, active sensing, and exploratory behaviors, the mechanisms underlying the motor production of sniffs are not known.

## Results

### Norepinephrine elicits sniffs *in vivo* and *in vitro*

Systemic administration of psychostimulants and catecholamines robustly produce stereotyped sniffing in rodents^32^ that has been attributed to central effects of norepinephrine (NE), a neurotransmitter that plays a crucial role in the sympathetic nervous system and in memory formation, attention, and arousal^33,34^. To determine whether NE acts at preBötC to promote sniffing, we microinjected NE (100 µM; final concentration: ∼10 µM) into preBötC of head-fixed awake behaving mice. We measured breathing using a nasal thermistor and tracked facial movements in video recordings using machine learning detection of keypoints with FaceMap (**Figure 1A**)^20,35^. During unilateral injection of saline vehicle into preBötC, regular, eupneic breaths occurring between 2-4 Hz (2.66 ± 0.51 Hz; n=7) were interspersed with short bouts of higher frequency (>4 Hz) breaths accompanied by larger amplitude nasal and whisker movements, which we called sniffs (**Figure 1B**)^9^. Even transient increases in respiratory frequency consisting of bouts of just two breaths above 4 Hz, were associated with significantly increased nose and whisker movements (nose, left whisker, right whisker x- and y-maximum displacement: F(2,12)=14.9-47.7, p=2×10^-6^-0.0006, eupnea vs. 2 bursts: p=4×10^-5^-0.03, eupnea vs. 3+ bursts: p=0.0003-0.008, 2 bursts vs 3+ bursts: p=0.001-0.007; n=7; **Figure 1C, Table S1**), suggesting that these brief bouts are also distinct from eupneic breathing. The fraction of sniff breaths relative to total breaths taken during the analysis period (sniff fraction) was 0.49 ± 0.24 (n=7). During unilateral injection of NE, basal eupneic frequency (2.76 ± 0.43 Hz) did not change significantly; however, an increase in the occurrence of sniff bouts, measured as a significant increase in the sniff fraction (0.65 ± 0.24), was observed (burst freq: t(6)=-1.9, p=0.4; sniff fraction: t(6)=-7.2, p=0.01; n=7; **Figure 1D-F, Supplementary Video S1**).

**Figure 1.**
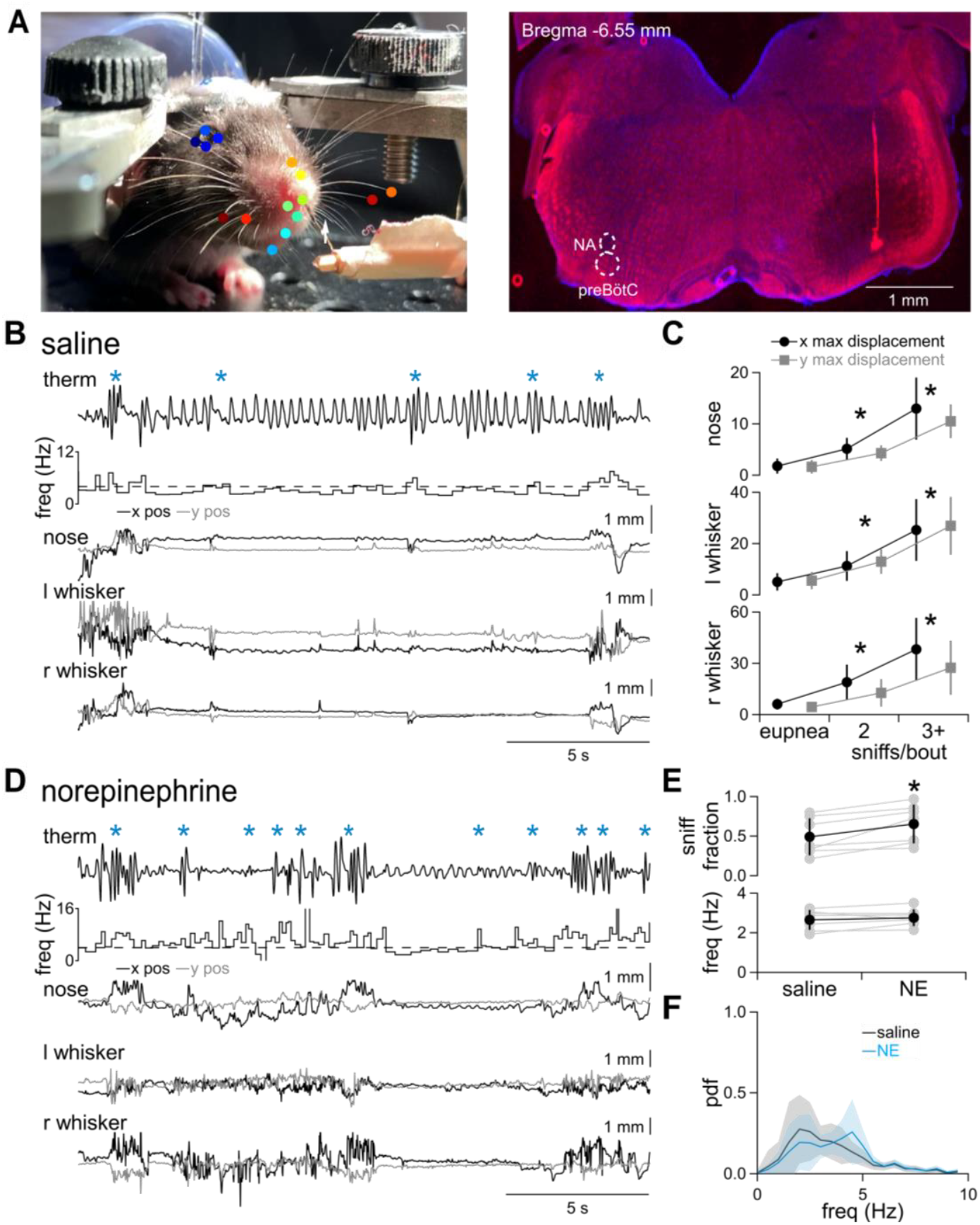
Norepinephrine increases sniffing *in vivo*. (**A**) *left*, Frame from video of head-fixed mouse with external nasal thermistor (white arrow) and Facemap-detected keypoints marked by colored dots. *right*, Coronal slice showing injection site marked by red fluorescent microspheres into preBötC. NA, nucleus ambiguus. (**B**) Representative traces of thermistor and x and y positions of FaceMap-detected keypoints for nose, left whisker (l whisker), and right whisker (r whisker) during unilateral injection of saline into preBötC. * indicate sniff bouts. Instantaneous respiratory frequency (freq) is shown below the thermistor recording. Dotted line at 4 Hz indicates our threshold for defining sniffs vs. eupnea. (**C**) Group data showing significantly increased nose and whisker movements associated with bouts of just 2 sniffs that are further increased for bouts of 3 or more (3+) sniffs (*, p<0.05 vs. eupnea; n=7). (**D**) Representative traces of thermistor and x and y positions of FaceMap-detected keypoints during unilateral injection of 100 µM norepinephrine (NE) into preBötC. * indicate sniff bouts. (**E**) Group data showing the effects of saline or NE on sniff fraction and eupneic breathing (<4 Hz) frequency. *, p<0.05, n=7. (**F**) Probability distribution function (pdf) of instantaneous frequency in saline or NE-injected animals. n=7.

To exclude NE effects on neighboring regions in the intact brainstem and determine the mechanisms underlying NE-induced sniff generation specific to preBötC, we applied NE to acute medullary slices containing a minimal breathing circuit that isolates the preBötC and generates rhythmic, respiratory-related activity^22,36^. We used extracellular field recordings to capture integrated rhythmic inspiratory population activity from preBötC (∫preBötC) and a physiological motor output, the hypoglossal motor nucleus or nerve (∫XII)^36^. In control (ctrl) conditions (9 mM K^+^/1.5 mM Ca^2+^ ACSF), ∫preBötC and ∫XII activity consisted of synchronous, 0.1-0.3 Hz inspiratory bursts that we suggest are eupneic and occasional longer duration, double peaked sigh bursts (**Figure 2A**)^23,36^. Consistent with previous studies^37-41^, bath application of NE (10 µM) increased burst duration with little effect on burst amplitude and frequency and did not produce sniff-like activity (amplitude: F(2,6)=0.3, p=0.7; duration: F(2,6)=102.6 p=2×10^-5^, ctrl vs. NE: p=7×10^-4^; frequency: F(2,6)=24, p=0.001, ctrl vs. NE: p=0.8; n=4; **Figure 2A-C**).

**Figure 2.**
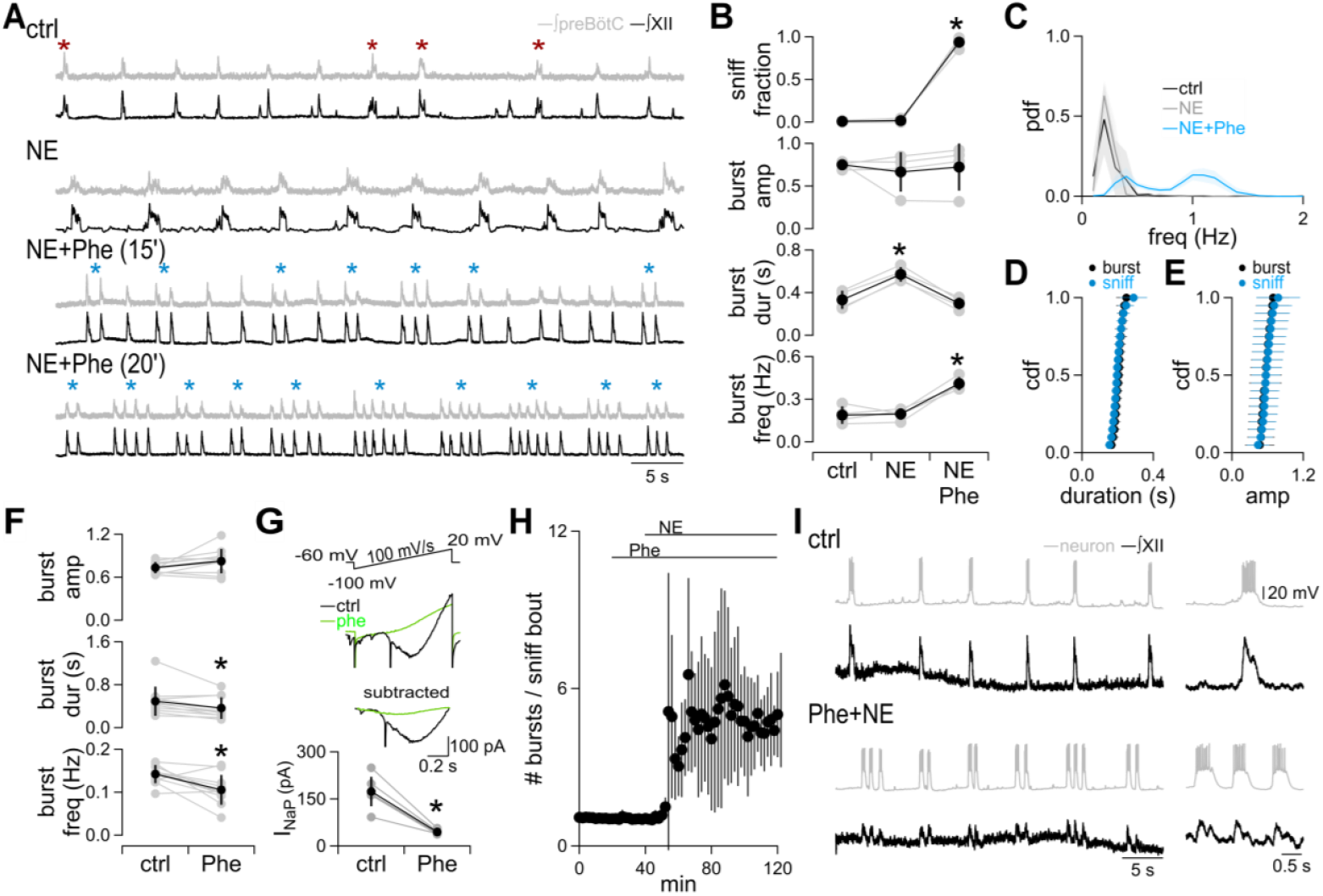
NE and Phe produce fictive sniffs. (**A**) Representative traces showing integrated preBötC (∫preBötC) and XII (∫XII) population recordings from rhythmic medullary slices in control (ctrl), NE (10 µM) and NE+Phenytoin (Phe: 100 µM) at two different time points (15’ and 20’) after beginning Phe application. * indicate fictive sighs. * indicate fictive sniffs. (**B**) Group data showing the effects of NE and NE+Phe on sniff fraction and burst amplitude (amp), duration (dur), and frequency (freq). *, p<0.05 vs ctrl, n=5. (**C**) Probability distribution function (pdf) of instantaneous burst frequency in control, NE, and NE+Phe. *, p<0.05, n = 5. (**D, E**) Cumulative distribution functions (cdf) of duration (D) and amplitude (E) of inspiratory bursts and bursts within sniff bouts, p>0.05, n=7. (**F**) Group data showing the effects of Phe on burst amp, dur, and freq. *, p<0.05, n=11. (**G**) *top*, I_NaP_ could be elicited with a 100 mV/s ramp that was blocked by Phe in whole-cell voltage clamp recordings from inspiratory preBötC neurons. I_NaP_ was isolated by subtracting a line fit to the linear portion of the current response. *bottom*, Group data showing a significant block of I_NaP_ by Phe. *, p<0.05, n=5. (**H**) Time course of effects of Phe and NE on the number of bursts in a sniff bout binned over 120 s. n = 11. (**I**) *left*, Representative traces of a whole cell current clamp recording of a preBötC inspiratory neuron and ∫XII activity in control and Phe+NE. *right*, The first burst is shown on an expanded time scale.

To generate high frequency breathing patterns like sniffing in a reduced slice preparation, we hypothesized that additional modulation of burst-generating mechanisms would be required and focused on I_NaP_, whose contributions *in vitro* and *in vivo* appear to differ. While I_NaP_ is important for shaping bursts and supporting inspiratory rhythmogenesis *in vitro*^22,42-44^, pharmacological or genetic knockdown of I_NaP_ in mouse models *in vivo* has only modest effects^42,45-47^. In humans, an I_NaP_ blocker, phenytoin (Phe), which is used therapeutically as an anti-epileptic, has not shown any central respiratory effects, and respiratory depression is not observed in cases of phenytoin overuse or overdose^46-49^. We therefore tested whether blocking I_NaP_ might be necessary to provide more *in vivo*-like conditions and elicit sniff-like activity *in vitro*^48^. Bath application of Phe (100 µM) after NE increased burst frequency, decreased burst duration, and did not change burst amplitude compared to NE alone (amplitude: F(2,6)=0.3, p=0.7; duration: F(2,6)=102.6, p=2×10^-5^, NE vs. Phe+NE: p=0.002; frequency: F(2,6)=24, p=0.001, NE vs. NE+Phe: p=0.005; n=4; **Figure 2A-C**), but the most striking effect was robust production of clusters of two or more bursts with ∼1 s interburst intervals (0.8-1.7 Hz) that were two to five times faster than baseline intervals (**Figure 2A-C**). Unlike sigh bursts, where activity persisted between the two peaks, bursts within these clusters were separated by quiescent periods (**Figure 2A**). We called these bouts fictive sniffs due to their resemblance with sniffs observed *in vivo* and *in situ* (**Figure 1B**)^7,50^.

To quantify this effect, we calculated a sniff fraction, i.e., the number of consecutive bursts with intervals <1.3 s divided by the total number of bursts. Adding Phe to NE significantly increased sniff fraction compared to control and NE alone (F(2,6)=557, p=2×10^-7^, ctrl vs. NE+Phe: p=0.0001, NE vs. NE+Phe: p=0.0002; n=4; **Figure 2A-C**). This “fast-slow” rhythm was evident in the bimodal distribution of interburst intervals in fictive sniff-generating conditions that was significantly different from the unimodal distributions in control or NE (ctrl vs NE+Phe: D(40)=0.4, p=2×10^-7^; NE vs NE+Phe: D(40)=0.5, p=1×10^-8^; n=4; **Figure 2C**). The bursts in these clusters were not significantly different from the slower inspiratory bursts in amplitude or duration (amplitude: D(70)=0.1, p=0.6; duration: D(70)=0.2, p=0.06; n=7; **Figure 2D, E)**.

To understand the effects of Phe that might contribute to fictive sniff generation, we applied Phe alone. Bath application of 100 µM Phe significantly reduced burst duration and frequency with no significant effect on burst amplitude (amplitude: t(10) = -4.1, p=0.07; duration: t(10)=5.9, p=0.01; frequency: t(10)=6.5, p=9×10^-3^; n=11; **Figure 2F**). To confirm that Phe blocks I_NaP_, we recorded inspiratory preBötC neurons in whole-cell voltage clamp using a Cs^+^ based internal with the non-specific K^+^ channel blocker tetraethylammonium^42^. We activated I_NaP_ in these neurons using a voltage ramp protocol (100 mV/s), which produced an inward current that activated around -20 mV, consistent with I_NaP_. A linear region was subtracted from the trace after fitting a line to obtain a peak I_NaP_ amplitude that measured 173.56 ± 46.85 mV (n=5). Bath application of 100 µM Phe significantly reduced I_NaP_ to 44.77 ± 6.43 mV (t(4)=5.7, p=2×10^−3^; n=5; **Figure 2G**), with the remaining current representing a Na^+^ window current not involved in bursting^42^.

In 100 µM Phe, NE (10 µM) increased burst frequency and did not change burst amplitude or duration compared to Phe (amplitude: F(2,16)=1.3, p=0.3, Phe vs. Phe+NE: p=0.6; duration: F(2,16)=7.7, p=0.005, Phe vs. Phe+NE: p=1; frequency: F(2,12)=37.2, p=7 x10^-6^, Phe vs. Phe+NE: p=0.0002; n=7; **Figure S1**). Similar to applying Phe after NE, adding NE to Phe significantly increased sniff fraction compared to control and Phe (F(2,12)=8548.4, ctrl vs. Phe+NE: p=4×10^-11^, Phe vs. Phe+NE: p=2×10^-11^; n=7; **Figure S1**) and produced a bimodal distribution of interburst intervals in fictive sniff-generating conditions that was significantly different from the unimodal distributions in control or Phe (ctrl vs Phe+NE: D(70)=0.5, p=2×10^-18^; Phe vs. Phe+NE: D(70)=0.5, p=6×10^-19^; n=7; **Figure S1**). The average number of consecutive high frequency sniff bursts, which we termed a sniff bout, in Phe and NE was 4.79 ± 0.69 (n=11) and could be stably generated for an hour (**Figure 2H**). NE also produced similar fictive sniff inspiratory patterns in the presence of riluzole, a more specific I_NaP_ blocker (**Figure S2**)^26,44,51^.

To determine whether fictive sniffs arose from the same population of preBötC neurons generating eupneic inspiratory activity, we performed whole cell patch clamp recordings. Inspiratory preBötC neurons generated bursts of activity synchronized with XII motor output in control and in Phe (**Figure 2I)**. Addition of NE led to fictive sniff activity in the same inspiratory preBötC neurons that occurred simultaneously with fictive sniffs recorded in XII output (n=9; **Figure 2I**).

The similarity in burst amplitude and duration and the resemblance of the frequency of fictive sniff bursts within sniff bouts to *in vivo* eupneic frequencies raises the possibility that these bouts are, in fact, eupneic. The frequency of eupneic rhythm *in vitro* and *in vivo* is sensitive to opioids and extracellular K^+36,52-55^. Adding the µ-opioid receptor agonist DAMGO (30 nM) or lowering extracellular K^+^ to 3 mM after Phe and NE did not reduce sniff fraction (F(5,20)=808; Phe+NE vs. 3 mM K^+^: p=0.5; Phe+NE vs. DAMGO: p=0.3; n=5; **Figure 3A, B**). Lowering extracellular K^+^ from 9 mM to 3 mM shifted the bimodal frequency distribution, significantly decreasing the inter-sniff bout frequency while increasing the intra-sniff bout frequency (pdf: D(50)=0.3, p=0.003; inter-sniff bout frequency: D(50)=0.6, p=1×10^-15^; intra-sniff bout frequency: D(50)=0.4, p=2×10^-8^; n=5; **Figure 3A-D**). DAMGO also significantly decreased the inter-sniff bout frequency but did not significantly change the bimodal frequency distribution or the intra-sniff bout interval (pdf: D(50)=0.1, p=0.4; inter-sniff bout frequency: D(50)=0.4, p=1×10^-6^; intra-sniff bout frequency: D(50)=0.1, p=0.7; n=5; **Figure 3A-C, E**). Thus, manipulations that depress eupneic rhythm either did not change or actually increased frequency of sniffs within bouts, pointing to a distinct mechanism governing sniff rhythm.

**Figure 3.**
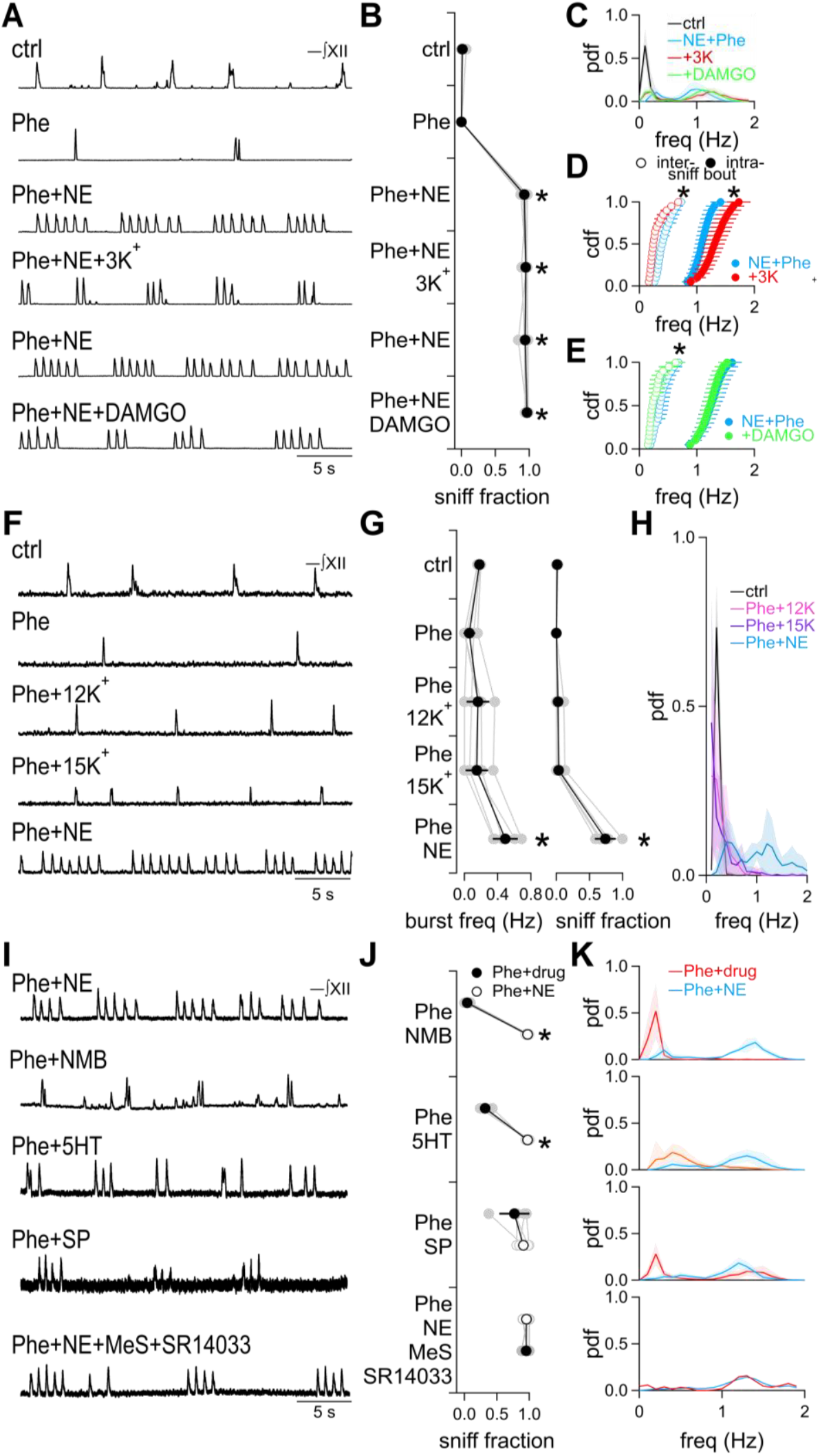
Fictive sniff rhythmogenesis is distinct from eupneic mechanisms and elicited by specific neuromodulators. (**A**) Representative ∫XII traces from a single slice experiment where fictive sniffs were elicited by Phe+NE, followed by lowering extracellular K^+^ to 3 mM, returning to 9 mM K^+^ and then bath application of the µ-opioid receptor antagonist DAMGO (30 nM). (**B**) Group data showing effects of 3 mM extracellular K^+^ and DAMGO on sniff fraction. *, p<0.05 vs. ctrl. n = 5. (**C**) Pdfs of instantaneous burst frequency in control, Phe+NE, Phe+NE+3 mM K^+^, and Phe+NE+DAMGO. N = 5. (**D, E**) Cdfs of instantaneous inter-sniff bout eupneic burst frequency (○) or intra-sniff bout frequency (●) in Phe+NE and either 3 mM K^+^ (D) or DAMGO (E). *, p<0.05, Phe+NE vs. 3 mM K^+^ or vs. DAMGO. n = 5. (**F**) Representative ∫XII traces from a single experiment showing the effects of raising extracellular K^+^ from 9 to 12 and 15 mM, and application of NE in Phe. (**G**) Group data showing the effects of raising extracellular K^+^ to 12 mM and 15 mM in Phe, and Phe+NE on sniff fraction and burst frequency. *, p<0.05 vs. ctrl. n = 5. (**H**) Pdfs of instantaneous burst frequency in control, Phe+12K^+^, Phe+15K^+^, and Phe+NE. n = 5. (**I**) Representative ∫XII traces from multiple experiments showing the effects of NE, neuromedin B (NMB; 30 nM), serotonin (5HT; 10 µM), Substance P (SP; 1 µM), and the serotonin and neurokinin 1 receptor antagonists methysergide (MeS; 10 µM) and SR14033 (10 µM) in Phe. (**J**) Group data showing effects of various neuromodulators (●) compared to NE (○) in Phe on sniff fraction. Upper condition was applied first. *, p<0.05 vs. NE+drug. n = 3-5. (**K**) Pdfs of instantaneous burst frequency of corresponding drugs in (J): NE or NMB (n = 4) (*top*), 5HT (n = 5) (*2*^*nd*^ *from top*), SP (n = 5) (*2*^*nd*^ *from bottom*), or NE+MeS+SR14033 (n = 3) (*bottom*) in Phe.

To test whether raising excitability independent of neuromodulation could also elicit sniffs, we raised extracellular K^+^ with I_NaP_ blocked. Raising extracellular K^+^ to 12 mM (12K^+^) and 15 mM (15K^+^) after Phe did not elicit sniffs (**Figure 3F-H**)^29,53^,56. Sniff fraction only increased significantly after adding NE in 9 mM K^+^ and Phe (F(4,16)=115.2, p=1×10^-11^, Phe vs. Phe+12K^+^: p=0.3, Phe vs. Phe+15K^+^: p=0.2, Phe+12K^+^ vs. Phe+NE: 0.0002, Phe+15K^+^ vs. Phe+NE: p=0.0001; n=5; **Figure 3F-H**). Decreasing K^+^ back to 9 mM and adding NE with Phe shifted the unimodal distribution of burst frequencies in 12 mM K^+^ and 15 mM K^+^ to a bimodal distribution consisting of faster frequencies (Phe+12K^+^ vs. Phe+NE: D(50)=0.3, p=5×10^-5^; Phe+15K^+^ vs. Phe+NE: D(50)=0.3, p=5×10^-5^; n=5; **Figure 3H**).

We then tested whether excitatory neuromodulators besides NE could also elicit fictive sniffs when I_NaP_ was blocked with Phe^57,58^. In Phe, the bombesin-like peptide neuromedin B (NMB; 30 nM) increased fictive sigh frequency compared to Phe alone (Phe vs. Phe+NMB: t(3) = -14.7, p=0.005; n=4), but did not significantly increase sniff fraction compared to control (F(3,9)=1903, p=6×10^-13^, ctrl vs. Phe+NMB: p=0.11; n=4), and the frequency distribution remained unimodal (**Figure 3I-K, S3**)^23^. After washing off NMB, subsequent application of NE in Phe in the same slices increased sniff fraction significantly and shifted the unimodal frequency distribution to a bimodal distribution (sniff fraction: F(3,9)=1903, p=6×10^-13^, Phe+NMB vs Phe+NE: p=2×10^-5^; pdf: Phe+NMB vs. Phe+NE: D(40)=0.6, p=7×10^-12^; n=4; **Figure 3I-K**). Bath-applied serotonin (5HT; 10 µM) in Phe increased sniff fraction compared to control, but NE in Phe produced significantly more sniffs than 5HT in the same slices (sniff fraction: F(3,12)=447, p=1×10^-12^, ctrl vs Phe+5HT: p=0.004, Phe+5HT vs Phe+NE: p=6×10^-5^; pdf: D(50)=0.1, p=0.8; n=5; **Figure 3I-K, S3**)^59,60^. Substance P (SP; 1 µM) produced robust fictive sniffs in Phe with sniff fraction significantly increased compared to control and not significantly different from a subsequent application of NE in Phe (sniff fraction: F(3,12)=61.1, p=2×10^-7^, ctrl vs Phe+SP: p=4×10^-3^, Phe+SP vs Phe+NE: p=0.3; pdf: D(50)=0.1, p=0.4; n=5; **Figure 3I-K, S3**)^60-62^. To determine whether NE activated serotonergic raphé neurons that also release SP, we blocked serotonin receptors with methysergide (10 µM) and neurokinin-1 receptors with SR14033 (10 µM) during NE application^60^. Co-application of the antagonists with NE in Phe did not significantly change the sniff fraction (sniff fraction: F(2,4)=270.3, p=5×10^-5^, Phe+NE vs Phe+NE+MeS+SR14033: p=1; pdf: D(30)=0.2, p=0.5; n=3; **Figure 3I-K**), suggesting that NE was capable of eliciting fictive sniffs directly, presumably by activating adrenergic receptors on preBötC neurons^37-41^.

### I_h_ underlies sniffs *in vitro* and *in vivo*

To determine the mechanisms by which NE induced rhythmic fictive sniffs in Phe, we tested the pacemaker conductances I_CAN_ and I_h_^26-28,42,63,64^ .Application of the I_CAN_ blocker 9-Phenanthrol (9-Phe; 100 µM) did not significantly affect frequency or amplitude but decreased duration (amplitude: F(2,8)=14.9, p=0.002, ctrl vs. 9-Phe: p=0.9; duration: F(2,8)=24.6, p=4×10^-4^, ctrl vs. 9-Phe: p=0.007; frequency: F(2,8)=16, p=0.002, ctrl vs. 9-Phe: p=0.6; n=5; **Figure 4A-C**)^65^. Blockade of I_CAN_ with 9-Phe had little effect on fictive sniffing as addition of Phe+NE in 9-Phe significantly increased sniff fraction (F(2,8)=57.9, p=2×10^-5^, 9-Phe vs. 9-Phe+Phe+NE: p=0.002; n=5; **Figure 4A-C**) and produced a bimodal distribution of frequencies reflecting robust fictive sniff generation (9-Phe vs. 9-Phe+Phe+NE: D(50)=0.6, p=1×10^-16^; n=5; **Figure 4A-C**).

**Figure 4.**
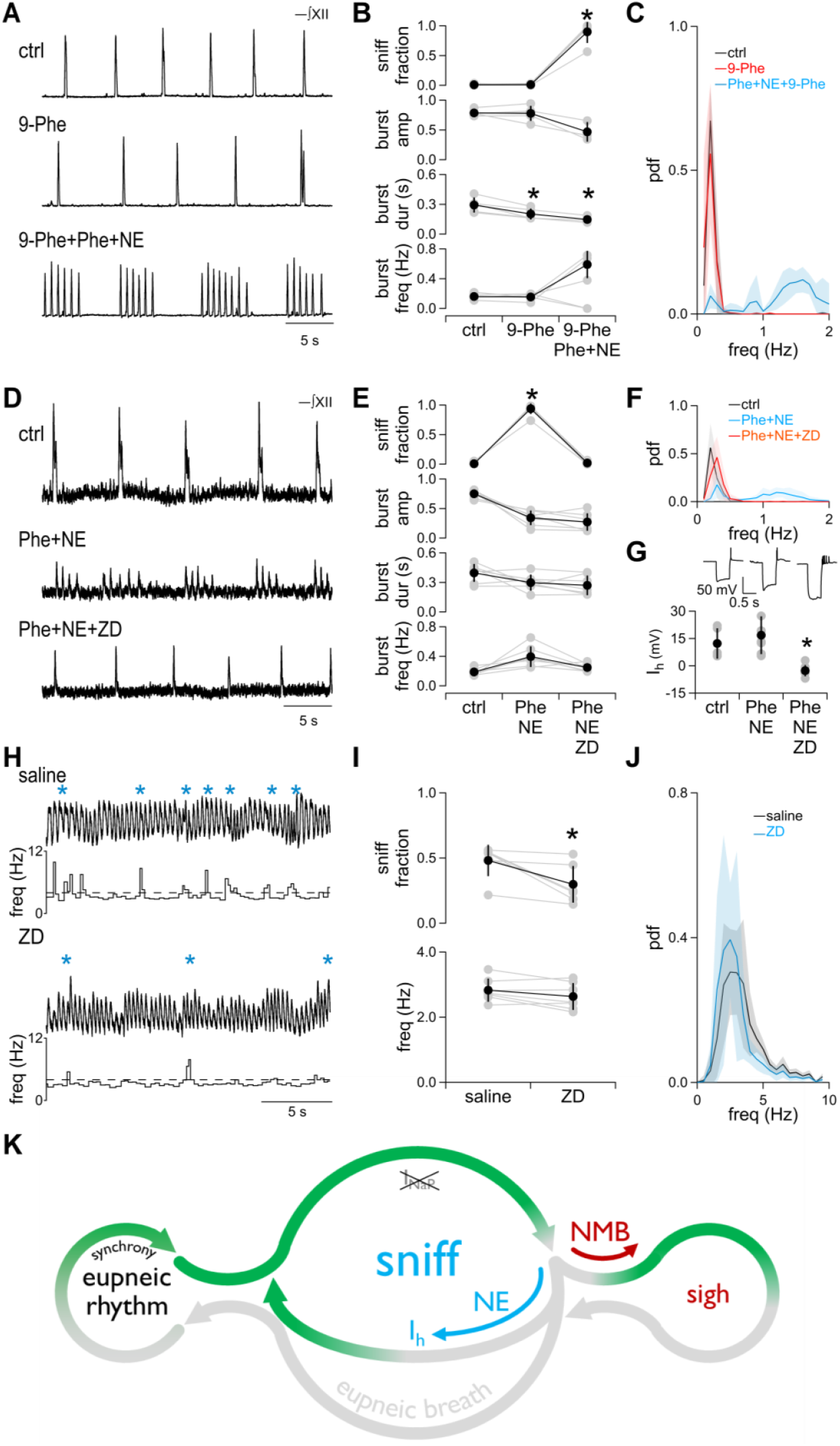
Fictive sniffs are dependent on I_h_. (**A**) Representative ∫XII traces from a single slice experiment showing the effects of I_CAN_ blocker 9-Phenanthrol (9-Phe; 100 µM). (**B**) Group data showing the effects of 9-Phe on sniff fraction and burst amp, dur, and freq. *, p<0.05 vs. ctrl. n = 5. (**C**) Pdfs of instantaneous burst frequency in control, Phe+NE, and Phe+NE+9-Phe. n = 5. (**D)** Representative ∫XII traces from a single slice experiment showing the effects of the I_h_ blocker ZD7288 (ZD; 30 µM). (**E**) Group data showing the effects of ZD on sniff fraction and burst amp, dur, and freq. *, p<0.05 vs. ctrl. n = 7. (**F**) Pdfs of instantaneous burst frequency in control, Phe+NE, and Phe+NE+ZD. n = 7. (**G**) Effects of ZD on I_h_ recorded in inspiratory preBötC neurons in whole cell current clamp mode. *Top*, Representative traces show the characteristic sag current attributed to I_h_ during a hyperpolarizing current pulse in control and Phe+NE that is abolished in ZD. *Bottom*, Group data showing I_h_ magnitude in control, Phe+NE, and Phe+NE+ZD. *, p<0.05. n = 10. (**H**) Representative traces of thermistor after bilateral injection of 100 nL of either saline (*top*) or 1 mM (final concentration 100 µM) ZD (*bottom*) into preBötC. * indicate sniff bouts. (**I**) Group data showing the effects of saline or ZD on sniff fraction and eupneic breathing (<4 Hz) frequency. *, p<0.05, n = 6. (**J**) Pdfs of instantaneous frequency in saline or ZD-injected animals. *, p<0.05. n = 6. (**K**) Model of preBötC eupnea and sniff generation, whereby conversion of eupneic bursts to sniffs depends on a norepinephrine activated, I_h_-dependent pacemaking mechanism.

In contrast, blockade of I_h_ with ZD7288 (ZD; 30 µM) abolished fictive sniffs while a slower eupneic rhythm remained (**Figure 4D-F**)^45,64^. ZD significantly reduced sniff fraction compared to Phe+NE to levels comparable to those in control conditions (F(2,12)=591, p=1×10^-12^, ZD vs. Phe+NE: p=1×10^-7^, ZD vs. ctrl: p=0.4; n=7; **Figure 4D-F**). ZD also produced a unimodal frequency distribution that was shifted to slower frequencies and overlapped with the unimodal distribution in control conditions (ZD vs. Phe+NE: D(70) = 0.5, p=4×10^-16^; ZD vs. ctrl: D(70) = 0.1, p=1.0; n=7; **Figure 4D-F**). Applying ZD before Phe+NE prevented generation of fictive sniffs (**Figure S4**). We measured I_h_ in inspiratory preBötC neurons recorded with whole cell patch clamp. 500 ms hyperpolarizing current injections produced a characteristic sag that is attributed to I_h_^63^. Phe and NE produced fictive sniffs and increased Ih in some neurons, but the overall effect was not significant (F(2,19)=11.6, p=5×10^-4^, ctrl vs. Phe+NE: p=0.1; n=10; **Figure 4G**), perhaps due to heterogeneity in preBötC I_h_ expression^63,66^. Subsequent bath application of ZD blocked both fictive sniffs and I_h_ (F(2,19)=11.6, p=5×10^-4^, ctrl vs. Phe+NE+ZD: p=0.001; n=10; **Figure 4G**). To test whether I_h_ contributes to sniffing *in vivo*, we injected ZD into preBötC in head-fixed awake behaving mice. Following bilateral injection of ZD (0.1 µL/side; final concentration: 100 µM), we observed significantly decreased sniff fraction with no significant effect on eupneic breathing frequency (burst freq: t(6)=-4.2, p=0.08; sniff fraction: t(6) = 7, p = 0.01; n=7; **Figure 4H-J**).

## Discussion

Our results demonstrate that sniffing is driven by a noradrenergic, I_h_-dependent pacemaker system that is conditionally activated during an ongoing eupneic rhythm governed by a distinct emergent process. In our model (**Figure 4K**), eupneic rhythmogenesis times the initiation of an active phase that can result in a single eupneic breath or, with NE present, bouts of higher frequency sniffs. We speculate that NE activates β-adrenergic receptors on preBötC neurons, which are coupled to G_s_ signaling pathways, to increase cAMP and positively modulate I _h_ ^67^. During each sniff, burst-driven Na^+^ and Ca^2+^ influx activating outward currents^68^, such as KCNQ^69^, may hyperpolarize neurons sufficient to activate the augmented I_h_, which is then able to quickly generate the next sniff.

Do the bouts of higher frequency inspiratory bursts in rhythmic medullary slices represent *in vitro* sniffs? While both fictive eupneic and sniff bursts *in vitro* are slower than the 1-2 Hz and 4-10 Hz eupneic and sniff rate in awake rodents, the relative increase in frequency, rapid transitions, and short sniff bouts are analogous^6,9^,20. Despite having similar frequencies, fictive sniffs are not likely to represent eupneic breathing since lowering extracellular K^+^ and applying opioids, which strongly depress eupneic rhythm^36,54^, did not affect or even increased intra-sniff bout frequency while prolonging inter-sniff bout intervals that we argue represent the emergent eupneic process. Additionally, the fictive sniffs we describe closely resemble sniffs evoked in anesthetized mice and odorant-evoked sniff-like activity in the more intact *in situ* rat nose-brainstem preparation^7,50^. We further show that *in vitro* and *in vivo* sniffs share sensitivity to NE and ZD when applied to preBötC. Therefore, sniff-like inspiratory patterns observed *in vitro* likely reflect physiological sniffs *in vivo*.

Eliciting sniffs *in vitro* requires I_NaP_ blockade. The raised excitability required for rhythmic activity in medullary slices may artificially augment I_NaP_ *in vitro* and prolong burst duration^22,36^,56. Increased Na^+^ and Ca^2+^ influx during longer bursts may amplify outward currents that extend a post-burst refractory period, preventing sniffs. Given the modest effects of pharmacological blockade or genetic knockdown of I_NaP_ *in vivo* in mice and humans^26,42^,45-47, we suggest that phasic I_NaP_ activation contributes minimally to inspiratory rhythm *in vivo* and that blocking I_NaP_ in slices brings *in vitro* conditions closer to the *in vivo* state.

Blockade of I_NaP_ reveals a novel effect of excitatory neuromodulators on preBötC. Neuromodulation expands the dynamic behavior in rhythmic networks^70^, and the breathing CPG is subject to extensive neuromodulation^57^. In contrast to increases in burst frequency or duration shown previously^40^, NE, with I_NaP_ blocked *in vitro*, altered preBötC dynamics from a single mode eupneic rhythm to a multimodal rhythm with fast and slow sniff and eupneic components. Noradrenergic locus coeruleus (LC) neurons project to preBötC and likely release NE during states where arousal or attention are increased^37^. SP and, more weakly, 5HT also increased sniff fraction with I_NaP_ blockade *in vitro*, pointing to possible additional contributions from these neuromodulator systems^60,62,71-73^. Consistent with the involvement of SP, photostimulation of parabrachial tachykinin-1 expressing neurons that encode the expression of several neuropeptides including SP and project to preBötC increased sniffing^19,74,75^.

Manipulations in awake, head-fixed mice support the role of NE in preBötC for sniff generation. Unilateral injection of NE into preBötC was sufficient to increase sniffing significantly without affecting eupneic breathing. While a limitation of drug injections *in vivo* is spread to neighboring regions, we argue that NE acts on preBötC neurons since the *in vivo* effects were consistent with our *in vitro* results, where preBötC is largely isolated. Nearby raphé neurons also present in the slice did not contribute significantly to the NE-induced increase in sniff fraction since blocking 5HT and NK1 receptors did not decrease fictive sniffs. While prior work showed that NE increases burst duration and frequency *in vitro* and increases sighing and stabilizes rhythm in anesthetized animals *in vivo*^39-41^, such effects may manifest differently when the animal is awake, highlighting the importance of validating results in unanesthetized animals and examining all aspects of breathing behavior, including coordinated orofacial movements^9,45^.

We suggest that NE activates I_h_ since ZD decreases sniffing both *in vitro* and *in vivo*. While ZD may also affect Na^+^ channels and synaptic transmission at concentrations that block I_h_^77,78^, we showed that ZD blocks I_h_ in slices, but did not affect eupneic rhythm, which depends on action potential firing and synaptic transmission, *in vitro* or *in vivo*, similar to previous work in anesthetized and awake mice^45,63.^ The involvement of I_h_ is further supported by the increased intra-sniff bout frequency when K^+^ was lowered, consistent with a Nernst-based hyperpolarization augmenting an I_h_-dependent mechanism^79-81^. ZD microdialysis in awake mice was previously shown to decrease breathing frequency when baseline breathing rate was high, perhaps congruent with a specific effect on fast breathing/sniffing, though only periods of regular breathing were analyzed^45^. The *in vivo* effects on sniffing we observed were more modest compared to *in vitro* experiments. The slow pharmacokinetics of ZD and poor tissue penetration of the lipophilic drug may have limited its effect^45,82^. Another possibility is that multiple mechanisms may contribute to sniff generation *in vivo*. These may be reflected in lower and higher frequency “resting” and “exploratory” sniffs that we find are associated with increasingly larger nose and whisker movements^83,84^.

The involvement of I_h_ and sympathetic-like noradrenergic regulation in sniffing places a cardiac-like pacemaker system in the brainstem that generates a biological rhythm for arousal and exploration as critical for survival as the heartbeat. Rapid shifts between eupneic breathing for gas exchange and sniff-like breaths for active olfactory sensation, arousal, and airway clearance enable humans and other mammals to switch respiratory function to meet the demands of the moment.

We suggest that during periods of arousal or attention, NE released from LC promotes sniff generation^37^. A reciprocal connection from preBötC Cdh9 neurons to LC may reinforce a state of arousal^85^ and, in cases of pathologically heightened arousal states, such as panic or anxiety disorder, mediate a feedback loop that underlies the rapid breathing that is a hallmark of a panic attack^17^. Conversely, deficits in LC function may occur in SIDS or CCHS and lead to a failure to generate sniff-like rescue breaths and an inability to rouse that can be fatal^4,15^. Uncovering this distinct mechanism for generating sniffing and fast sniff-like breathing enhances our understanding of respiratory and olfactory sensorimotor circuitry underlying a suite of active sensing and exploratory behaviors and reveals how multiple respiratory-related rhythms can be generated in the brainstem to shape breathing, protect the airway, initiate arousal, and coordinate orofacial motor activity.

## Funding

National Institutes of Health grant R01NS097492 (KK)

DePaul-RFUMS Biomedical Innovations grant (KK)

Chicago Biomedical Consortium (KK)

National Institutes of Health grant R56MH130488 (EJH)

Alfred P. Sloan Fellowship (EJH)

DePaul-RFUMS Biomedical Innovations grant (EJH)

## Author contributions

Conceptualization: BT, KK

Methodology: BT, EJH, KK

Investigation: BT, YL, JS, MCA

Visualization: BT, KK

Funding acquisition: KK, EJH

Project administration: KK

Supervision: EJH, KK

Writing – original draft: KK

Writing – review & editing: BT, JS, EJH, KK

## Supplementary Materials

Figs. S1 to S4

Table S1

Video S1

## Materials and Methods

### Animals

The Institutional Animal Care and Use Committee at Rosalind Franklin University of Medicine and Science approved all the protocols in alignment with the guidelines of the National Research Council and policies of the Office of Laboratory Animal Welfare (National Institutes of Health). For *in vivo* experiments, adult (>P56) C57BL/6J wild-type mice (JAX 000664; 9 female, 5 male) were used. For *in vitro* experiments, we used P0-6 C57BL/6 or transgenic reporter mice on C57BL/6 backgrounds with a wild type phenotype, e.g., GlyT2^EGFP^ (JAX 038516), Ai14 (JAX 007914), or other reporter mice, of either sex. Maximum care was taken to minimize the pain and discomfort of animals.

### Stereotaxic surgery

Stereotaxic surgeries were performed under general anesthesia (1.5 - 2% isoflurane) under aseptic conditions to implant a head-fixation bar, create microcraniotomies above the preBötC, and label the preBötC injection target.

A stainless steel head-fixation bar was affixed to the skull ∼1 mm posterior to bregma, aligned parallel to the horizontal plane of the stereotaxic frame^86^. Then, bilateral micro-craniotomies were made above the preBötC (AP: -6.59 mm, ML: ±1.4 mm from bregma). To virally label the preBötC, a glass micropipette (∼25 µm tip diameter) loaded with pAAV-hSyn-EGFP (Addgene 50465-AAV1; titer: 2.71×10^12^ GC/mL) was lowered 4.72 mm below the dura through the microcraniotomy and 100 nL of virus was injected at 25 nL/min. After waiting ∼10 minutes for viral diffusion, the pipette was retracted.

Because the head-fixation bar was aligned to the stereotaxic coordinate frame, subsequent drug injections could be repeatedly targeted to the preBötC by securing the mouse in the head-fixation apparatus and lowering pipettes through the same microcraniotomies to the preBötC depth. To validate that future injections would reach the preBötC, mice were transferred from the stereotaxic frame to the head-bar fixation apparatus, and a second viral tracer (pENN.AAV.hSyn.TurboRFP.WPRE.RBG; Addgene 105552-AAV1; titer: 2.14×10^12^ GC/mL) was injected at the same depth (100 nL).

After this second injection, the skull was covered with dental acrylic (1530 BLK; Lang dental), leaving the craniotomies exposed. The craniotomies were then sealed with silicone sealant (World Precision Instruments KWIK-CAST) to allow repeated access during later drug injection sessions. During the ∼3-day postoperative recovery period, mice received daily subcutaneous meloxicam (1 mg / kg). General health and postoperative discomfort were monitored using body condition scoring and the mouse grimace scale.

### Drug injection

For each drug injection session, the mouse’s head was secured in the head-bar fixation apparatus, and the silicone sealant was removed from the craniotomies. A micropipette loaded with one of three treatments was advanced into the preBötC: 0.9% saline vehicle (control), norepinephrine (100 µM; final concentration 10 µM), or ZD 7288 (1 mM; final concentration 100 µM) injected at 10X to achieve the desired final concentration^87^. All injections were delivered at 25 nL/min. During each injection, the craniotomies were kept covered with 0.9% saline to prevent tissue drying.

In each session, breathing, sniffing, and general body movements were monitored and recorded using a video camera (FLIR CM3-U3-13Y3M-CS at 30 fps or Apple iPhone 14 camera at 60 fps) and a NTC thermistor (GAG22K7MCD419, TE Connectivity) connected to a TC-344C temperature controller (Warner Instruments). Thermistor signals were digitized at 1 kHz using a MiniDigi (Molecular Devices) and acquired using pClamp 9 (Molecular Devices). Thermistor and video signals were synchronized via a simultaneous auditory cue with an electronic pulse. For within-subject comparisons, each mouse received the different treatments on separate days over a period of 5-10 days. After injections were complete, pipettes were left in place for 10 mins to prevent backflow. After 10 mins, the pipettes were withdrawn, and the craniotomies were covered with silicone sealant. For NE, the analysis period occurred during injections on an open stereotaxic apparatus. Due to the slow pharmacokinetics of ZD, after the craniotomies were covered with the silicone sealant, mice were moved to an enclosed, darkened chamber, where breathing and facial movements were recorded. The analysis period for ZD occurred 15-35 minutes after bilateral injection of ZD was complete.

After the final drug injection session, 100 nL of latex beads (Sigma L3280) or Fast Blue (Polysciences) was injected into the preBötC under the same head-bar fixation condition. These bead/dye injections were used to assess the accuracy and consistency of targeting across sessions by comparing their localization to the viral expression sites labeled earlier.

### Perfusion and histology

Mice were sedated with a ketamine/xylazine mixture (100 mg / 20 mg / kg) and perfused transcardially with PBS followed by 4% paraformaldehyde (PFA). Brains were extracted, post-fixed in 4% PFA overnight, and then transferred to 30% sucrose for ∼48 hours. Coronal brain sections (50 µm) were prepared using a sliding microtome (SM2000R, Leica Microsystems) and imaged with a slide scanning fluorescence microscope (Thunder Imager 3D tissue, Leica Microsystems) to verify drug injection sites.

### *In vitro* slice preparation and electrophysiology

P0-P6 mice of either sex were anesthetized by inhalation of isoflurane, and the brainstem was quickly isolated in ACSF containing the following (in mM): 124 NaCl, 3 KCl, 1.5 CaCl_2_, 1 MgSO_4_, 25 NaHCO_3_, 0.5 NaH_2_PO_4_, and 30 D-glucose, equilibrated with 95% O_2_, and 5% CO_2_, 27-29°C, pH 7.4. Rhythmically active transverse medullary slices (550-650 µm thick) from brainstem containing preBötC, respiratory premotor neurons, and XII motoneurons were prepared as previously described^88^. To capture preBötC at the rostral surface, the 550-650 µm thick slice was cut after seeing nucleus ambiguus.

Slices with rostral side up were then stabilized in a ∼2 mL recording chamber under a fixed-stage upright microscope (Zeiss AxioExaminer Z1). To generate a robust and stable burst rhythm in the medullary slices, extracellular K^+^ was raised to 9 mM and extracellular Ca^2+^ was maintained at 1.5 mM in the recording ACSF. Slices were perfused with 27-29°C recording ACSF at 2.5-3 mL/min and allowed to equilibrate for at least 30 minutes to reach a steady-state in frequency and magnitude of XII and preBötC activity. To record respiratory-related activity, suction electrodes (∼ 50 µm tip size) were placed on preBötC and XII motor nucleus or XII nerve roots to record population activity, reflecting suprathreshold action potential (AP) firing. Signals were amplified in either a Model 1700 (A-M Systems) or Multiclamp 700B (Molecular Devices) amplifier, filtered at 2-4 kHz and digitized at 10 kHz. Integration was performed with a Paynter filter with a 20-100 ms time constant (Bak Electronics). To acquire continuous data, analog-to-digital converter (BNC-2090, National Instruments) and IgorPro (Wavemetrics) were used. Digitized data were later analyzed offline in custom procedures written in Igor Pro (Wavemetrics).

### Whole-cell patch clamp electrophysiology

Whole-cell patch-clamp recordings were obtained from preBötC inspiratory neurons. Neurons were visualized by infrared-enhanced differential interference contrast (IR-DIC) video microscopy with a 20X objective (Zeiss). Patch pipettes had a tip resistance of 4-6 MΩ and were fabricated using a Flaming-Brown micropipette puller with a three-stage custom program (P-87, Sutter Instruments). Whole-cell patch clamp was performed using a MultiClamp 700B amplifier (Molecular Devices). Signal was then filtered at 2-4 kHz and digitized at 10 kHz. To acquire continuous data, analog-to-digital converter (BNC-2090, National Instruments) and IgorPro (Wavemetrics) were used. To record persistent sodium current (I_NaP_) activity from preBötC inspiratory neurons, the patch pipette solution contained tetraethylammonium chloride (TEA-Cl) and Cs^+^ to minimize K^+^ currents, and had the following composition (in mM): 110 CsCl, 5 NaCl, 20 TEA-Cl, 10 EGTA, 10 HEPES, 1.2 CaCl_2_, 2 Mg-ATP, and 0.3 Na_3_-GTP^42^. To activate I_NaP_ and minimize activation of transient sodium current, a command ramp protocol of 100 mV/s speed from -100 mV to +20 mV with a baseline holding potential of -60 mV in voltage-clamp mode was utilized^42^. To record the presence of I_h_ in preBötC inspiratory neurons, hyperpolarizing current pulses 500 ms in duration were given in whole cell current clamp from a membrane potential of -60 mV to produce a hyperpolarizing response of -40 to -50 mV with a depolarizing sag. The internal solution had the following composition (in mM): 140 Kgluconate, 5 NaCl, 0.1 EGTA, 10 HEPES, 1 MgCl_2_, 2 Mg-ATP, 0.3 Na_3_-GTP^66^. Analysis of traces was performed offline using custom software written for IgorPro.

### Pharmacological reagents

Pharmacological reagents were prepared from stock solution and diluted to their final concentration. Phenytoin sodium was obtained from Spectrum Chemical Mfg., Corp. Riluzole, 9-Phenanthrol, DL-Norepinephrine hydrochloride, DAMGO, Substance P, Serotonin Hydrochloride, and Methysergide maleate salt were obtained from Sigma-Aldrich. Neuromedin B and SR 14033 were obtained from RnD Systems. ZD7288 was obtained from Hello Bio.

### Data analysis and statistics

Thermistor recordings were preprocessed in IgorPro by applying a smoothing filter with a time constant of 5 ms. This low pass filtered signal was then subtracted from the raw signal to reduce slow drift in temperature unrelated to respiratory airflow^89^. Breaths were semi-automatically detected using custom procedures written in IgorPro based on amplitude thresholds and confirmed visually by the experimenter.

60 fps video files in MOV format were converted to MP4 using VLC media player (VideoLan). Videos were imported into FaceMap^35^, and keypoints for eye front, eye top, eye back, eye bottom, chin, mouth, nose bottom, right nares, nose tip, nose top, nose bridge, two left whiskers and two right whiskers were selected. 25 frames from each video were used to refine the base model, and keypoints were evaluated for accuracy. Videos were then processed in FaceMap that was run in a Google CoLab environment. X- and y-positions for keypoints were measured, and a confidence measure was calculated. Timepoints when the confidence measure fell below 0.7 were blanked. To mitigate keypoint jitter, the five nose keypoints were averaged together to generate a nose trace. Similarly the two left whiskers were averaged together to get a left whisker trace, and the two right whiskers were averaged together to obtain a right whisker trace. Traces were manually aligned with the thermistor recording using the auditory-electronic synchronization signal.

Bursts of activity were detected from ∫XII output or ∫preBötC population activity using custom procedures written in IgorPro based on slope and amplitude thresholds and confirmed visually by the experimenter. Burst amplitude, duration, and frequency were measured using custom software written in IgorPro.

Following detection, bursts of activity were categorized as bursts, sighs, and sniffs based on interval and duration. Sighs were defined as two-peaked bursts with an interval between the two peaks <0.6 s and/or duration > 0.55 s. Fictive sniff bouts were consecutive bursts of activity >0.6s and <1.3 s and were defined as such due to their resemblance with sniffs observed *in vivo* and *in situ*^7,50^. Activity that did not fall into either category were defined as bursts. All bursts, sighs, and sniffs were visually inspected to verify categorization. Average burst, sniff, and sigh parameters were analyzed over a 10 minute period when rhythm had stabilized after drug application.

To isolate I_NaP_, a line was fit to the linear region at hyperpolarized potentials in the voltage-clamp ramp. This fitted line was then subtracted from the total current trace, revealing I_NaP_. The peak I_NaP_ amplitude was measured by subtracting the peak inward current from the baseline. The baseline and the peak inward current were calculated over a 20 ms period.

To measure the magnitude of depolarizing sag potentials by I_h_ activation in response to a hyperpolarizing current injection, the amplitude was quantified by the difference between the peak hyperpolarization and the steady-state membrane potential during the current step.

In group data, gray circles represent measures from individual experiments. Data are presented as mean ± SD, and statistical significance was set at a minimum of *p* < 0.05. Data were tested for normality using the Junque-Bera test. For normally-distributed data, parametric tests for repeated measures of two groups (paired Student’s t-test) or more than two groups (Two-way repeated measures ANOVA) were used. Post-hoc paired t-tests with Bonferroni-Holm correction were used to determine pairwise differences. When some samples produced no bursts in certain conditions, preventing the use of repeated measures, one-way ANOVA was used with Tukey post-hoc tests. Cumulative distribution functions (cdfs) were calculated after sorting and binning data into 20 bins with each bin containing 5% of the data points. Similarly, probability distribution functions (pdfs) were calculated after sorting and binning data into 20 bins with each having a width of 0.1 Hz. The histogram was normalized, so that the integral of pdf was 1. Kolmogorov-Smirnov tests were used for non-parametric comparisons of distributions.

