## Supplementary Materials for "A noradrenergic I_h_-dependent pacemaker system drives sniffing"

**Supplementary Materials include:**

Figs. S1 to S4
Table S1
Video S1

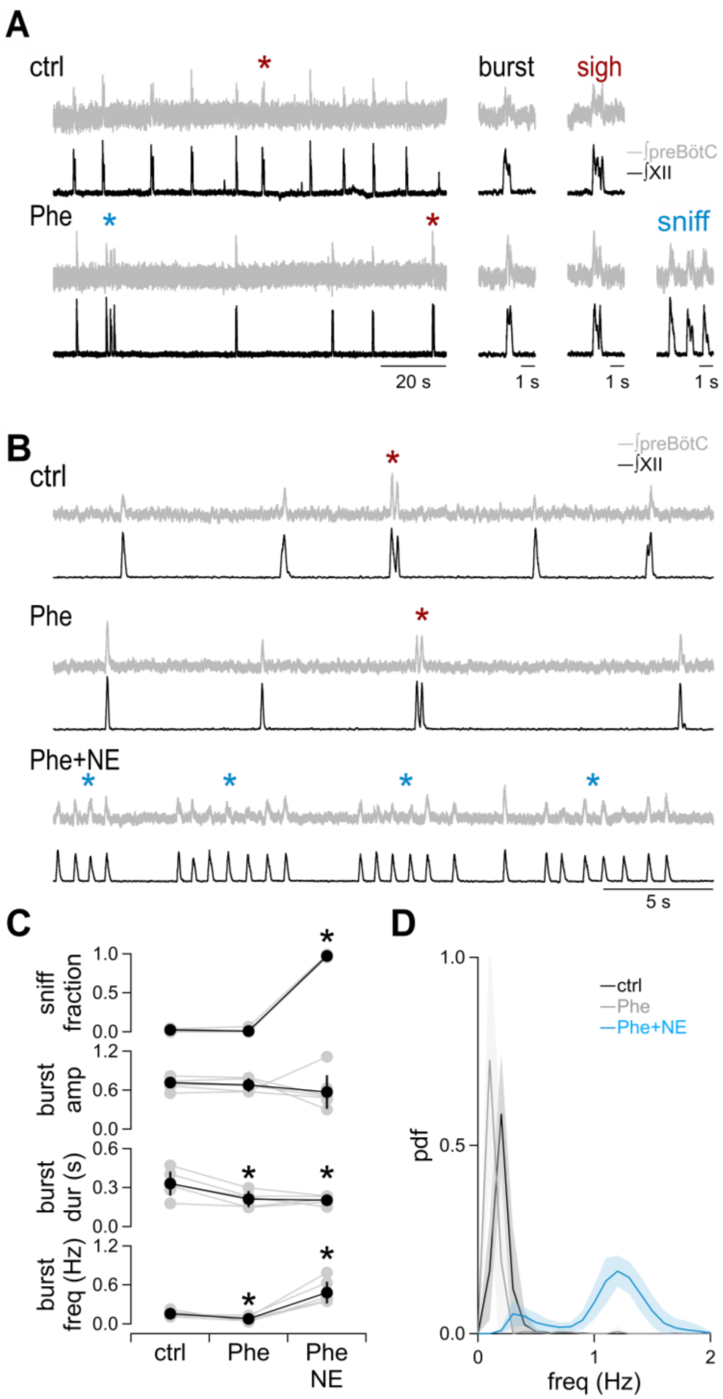


**Figure S1. Phe and NE produce fictive sniffs.**
(**A**) Representative traces showing integrated preBötC (∫preBötC) and XII (∫XII) population recordings from rhythmic medullary slices in control (ctrl) and phenytoin (Phe; 100 µM). Inspiratory-related activity in control consists of single-peaked burst and double-peaked sighs (*). In Phe, an occasional bout of higher frequency bursts, which we termed fictive sniffs (*), was observed. (**B**) Representative ∫preBötC and ∫XII traces in control, Phe, and Phe and norepinephrine (NE; 10 µM) together (Phe+NE). (**C**) Group data showing the effects of Phe and Phe+NE on sniff

fraction and burst amplitude (amp), duration (dur), and frequency (freq) (sniff fraction: F(2,12)=8548.4, ctrl vs. Phe+NE: p=4x10$^{-11}$, Phe vs Phe+NE: p=2x10$^{-11}$; amplitude: F(2,16)=1.3, p=0.3; duration: F(2,16)=7.7, p=0.005, ctrl vs Phe: p=0.02, ctrl vs Phe+NE: p=0.006, Phe vs. Phe+NE: p=1; frequency: F(2,12)=37.2, p=7x10$^{-6}$, ctrl vs Phe: p=0.008, ctrl vs Phe+NE: p=0.003, Phe vs. Phe+NE: p=0.0002; n=7). *, p<0.05. (**D**). Probability distribution function (pdf) of instantaneous burst frequency in control, Phe, and Phe+NE (ctrl vs Phe: D(70)=0.2, p=0.03; ctrl vs Phe+NE: D(70)=0.5, p=2x10$^{-18}$; Phe vs Phe+NE: D(70)=0.5, p=6x10$^{-19}$; n=7). *, p<0.05.

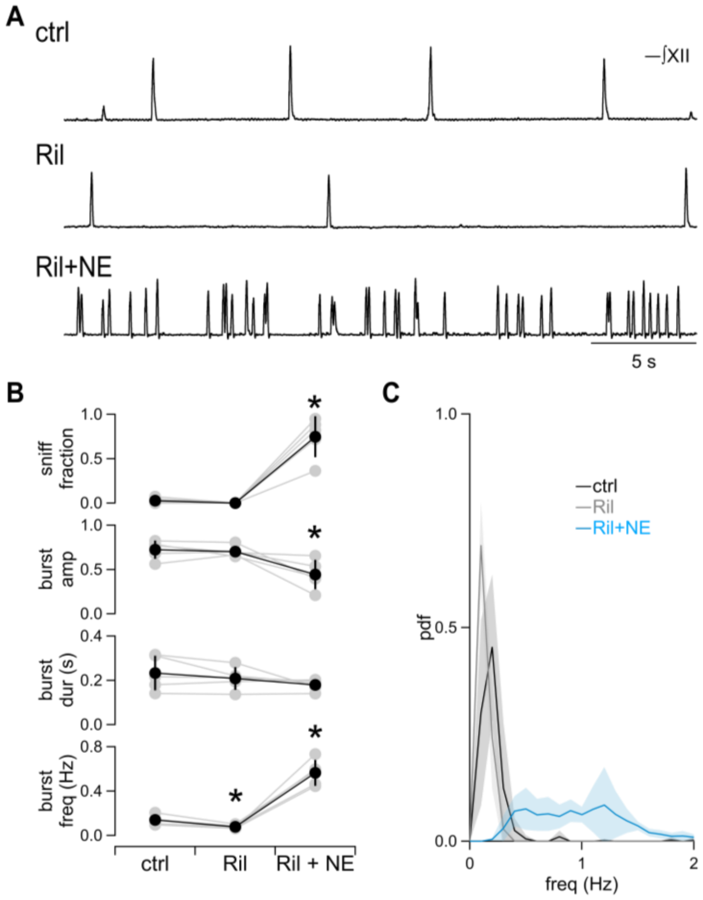


**Figure S2. The $I_{NaP}$ blocker riluzole and NE produce fictive sniffs.**
(**A**) Representative ∫XII traces from a single slice experiment showing the effects of riluzole (Ril; 10 µM) and Ril+NE (10 µM). (**B**) Group data showing the effects of Ril and Ril+NE on sniff fraction and burst amp, dur, and freq (sniff fraction: $F(2,8)=52$, $p=3x10^{-5}$, ctrl vs. Ril: $p=0.1$, ctrl vs. Ril+NE: $p=0.002$; Ril vs. Ril+NE: $p=0.002$; amplitude: $F(2,8)=12.7$, $p=0.003$, ctrl vs. Ril: $p=0.6$, ctrl vs. Ril+NE: $p=0.01$, Ril vs Ril+NE: $p=0.03$; duration: $F(2,8)=2.2$, $p=0.2$; frequency: $F(2,8)=87$, $p=4x10^{-6}$ , ctrl vs. Ril: $p=0.01$, ctrl vs. Ril+NE: $p=7x10^{-4}$, Ril vs. Ril+NE: $p=0.0007$; $n=5$). (**C**) Pdfs of instantaneous burst frequency in control, Ril, and Ril+NE (ctrl vs. Ril: $D(50)=0.1$, $p=0.3$, ctrl vs. Ril+NE: $D(50)=0.7$, $p=1x10^{-20}$, Ril vs. Ril+NE: $D(50)=0.8$, $p=3x10^{-29}$, $n = 5$). *, $p<0.05$.

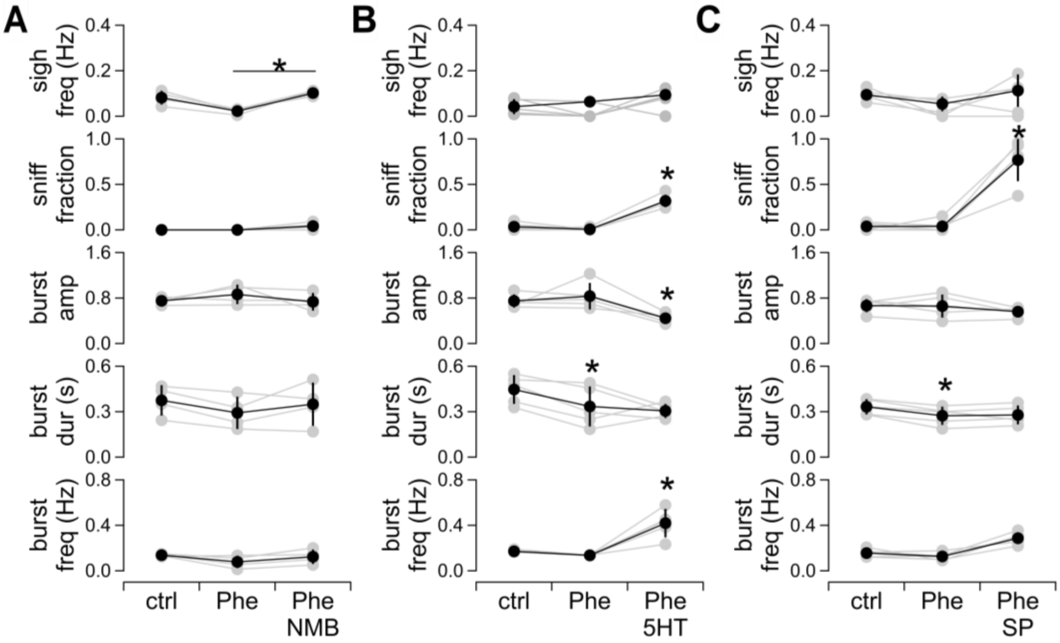


**Figure S3. Effects of neuromodulators in Phe.**
(**A**) Group data showing the effects of Phe (100 µM) and Phe+NMB (30 nM) on sniff fraction, sigh frequency, and burst amp, dur, and freq (sigh frequency: F(2,6)=17.7, p=0.003, Phe vs Phe+NMB: p=0.005; sniff fraction: F(2,6)=5, p=0.05; amplitude: F(2,6)=1.2, p=0.4; duration: F(2,6)=2.4, p=0.2; frequency: F(2,6)=1.9, p=0.2; n=4). (**B**) Group data showing the effects of Phe and Phe+serotonin (5HT; 10 µM) on sniff fraction, sigh frequency, and burst amp, dur, and freq (sigh frequency: F(2,8)=2.6, p=0.1; sniff fraction: F(2,8)=52.5, $p=3x10^{-5}$, ctrl vs Phe+Ser: p=0.004, Phe vs Phe+Ser: p=0.0002; amplitude: F(2.8)=9.5, p=0.008, ctrl vs Phe+Ser: p=0.01, Phe vs Phe+Ser: p=0.01; duration: F(2,8)=5.1, p=0.04, ctrl vs Phe: p=0.01; frequency: F(2,8)=34.3, p=0.0001, ctrl vs Phe+Ser: p=0.005, Phe vs Phe+Ser: p=0.003; n=5). (**C**) Group data showing the effects of Phe and Phe+SP (1 µM) on sniff fraction, sigh frequency, and burst amp, dur, and freq (sigh frequency: F(2,8)=2.1, p=0.2; sniff fraction: F(2,8)=40.2, $p=7x10^{-5}$, ctrl vs Phe+SP: p=0.003, Phe vs Phe+SP: p=0.002; amplitude: F(2,8)=1.9, p=0.2; duration: F(2,8)=6.9, p=0.02, ctrl vs Phe: p=0.007; frequency: F(2,8)=0.4, p=0.7; n=5). *, p<0.05.

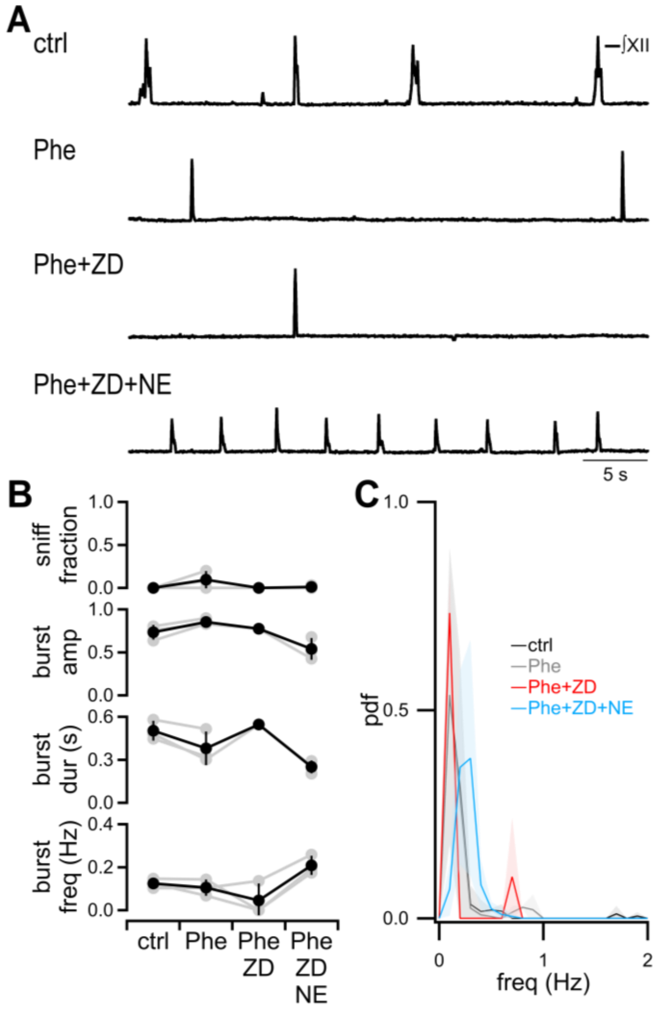


**Figure S4. $I_h$ blockade prevents NE from producing fictive sniffs.**
(**A**) Representative ∫XII traces from a single slice experiment showing the effects of Phe (100 µM), Phe+ZD7288 (ZD; 30 µM) and Phe+ZD+NE (10 µM). (**B**) Group data showing the effects of Phe, Phe+ZD, and Phe+ZD+NE on sniff fraction and burst amp, dur, and freq (sniff fraction: F(3,6)=2.7, p=0.1; amplitude: F(3,6)=6.1, p=0.03; duration: F(3,6)=5.9, p=0.03; frequency: F(3,6)=5.8, p=0.03; n = 3). (**C**) Pdfs of instantaneous burst frequency in control, Phe, Phe+ZD, and Phe+ZD+NE (ctrl vs. Phe+ZD+NE: D(30)=0.05, p=1.0, Phe vs. Phe+ZD+NE: D(30)=0.08, p=1.0; Phe+ZD vs. Phe+ZD+NE: D(30)=0.2, p=0.1; n = 3). *, p<0.05.

| | x max displacement | | | | | y max displacement | | | | |
|---|---|---|---|---|---|---|---|---|---|---|
| | F(2,12) | F(2,12) p-value | eupnea vs 2 sniffs p-value | eupnea vs 3+ sniffs p-value | 2 sniffs vs 3+ sniffs p-value | F(2,12) | F(2,12) p-value | eupnea vs 2 sniffs p-value | eupnea vs 3+ sniffs p-value | 2 sniffs vs 3+ sniffs p-value |
| **nose** | 22.9 | $8\text{x}10^{-5}$ | 0.003 | 0.003 | 0.004 | 47.7 | $2\text{x}10^{-6}$ | $4\text{x}10^{-5}$ | 0.0003 | 0.001 |
| **l whisker** | 16.9 | 0.0003 | 0.02 | 0.005 | 0.007 | 15.3 | $5\text{x}10^{-5}$ | 0.007 | 0.002 | 0.002 |
| **r whisker** | 18.6 | 0.0002 | 0.02 | 0.005 | 0.002 | 14.9 | 0.0006 | 0.03 | 0.008 | 0.004 |

**Table S1. Statistics for Figure 1C**

F statistics and p-values for data presented in Figure 1C.

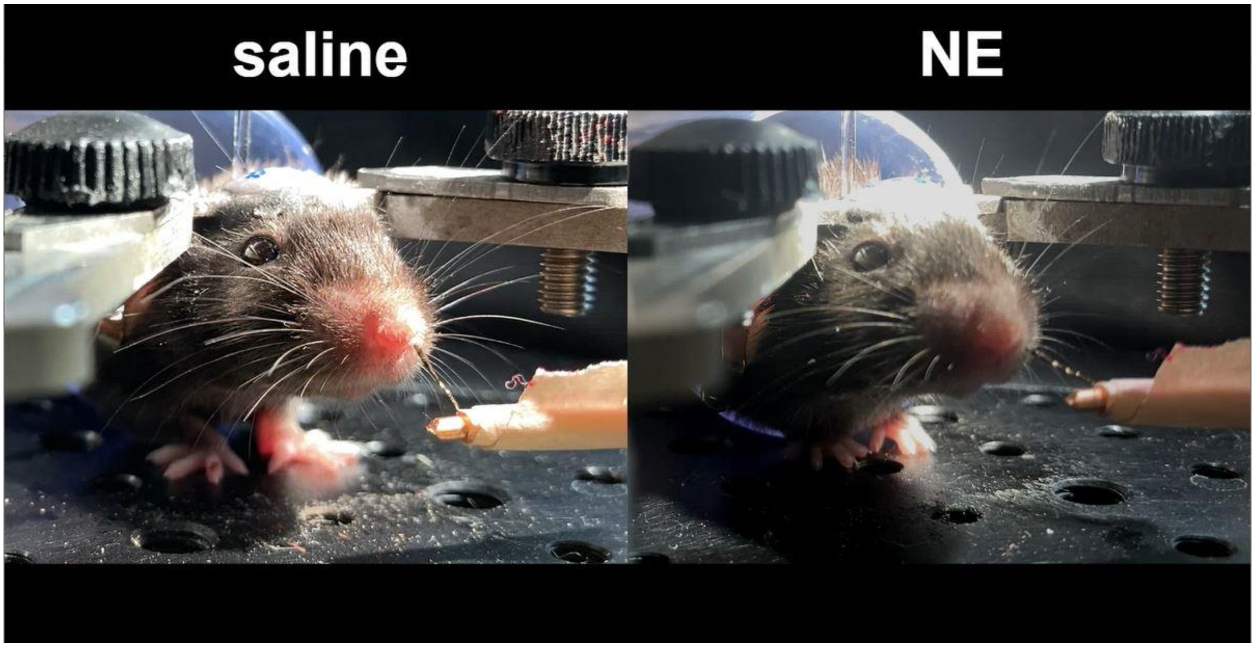


**Video S1. NE increases frequency of sniff bouts.**
*Left*, Head-fixed awake mouse during saline injection into preBötC. *Right*, Head-fixed awake mouse during NE injection into preBötC.
